# Hyaluronic acid-functionalized gelatin hydrogels reveal extracellular matrix signals temper the efficacy of erlotinib against patient-derived glioblastoma specimens

**DOI:** 10.1101/556324

**Authors:** Sara Pedron, Gabrielle L. Wolter, Jee-Wei E. Chen, Sarah E. Laken, Jann N. Sarkaria, Brendan A. C. Harley

## Abstract

Therapeutic options to treat primary glioblastoma (GBM) tumors are scarce. GBM tumors with epidermal growth factor receptor (EGFR) mutations, in particular a constitutively active EGFRvIII mutant, have extremely poor clinical outcomes. GBM tumors with concurrent EGFR amplification and active phosphatase and tensin homolog (PTEN) are sensitive to the tyrosine kinase inhibitor erlotinib, but the effect is not durable. A persistent challenge to improved treatment is the poorly understood role of cellular, metabolic, and biophysical signals from the GBM tumor microenvironment on therapeutic efficacy and acquired resistance. The intractable nature of studying GBM cell in vivo motivates tissue engineering approaches to replicate aspects of the complex GBM tumor microenvironment. Here, we profile the effect of erlotinib on two patient-derived GBM specimens: EGFR+ GBM12 and EGFRvIII GBM6. We use a three-dimensional gelatin hydrogel to present brain-mimetic hyaluronic acid (HA) and evaluate the coordinated influence of extracellular matrix signals and EGFR mutation status on GBM cell migration, survival and proliferation, as well as signaling pathway activation in response to cyclic erlotinib exposure. Comparable to results observed in vivo for xenograft tumors, erlotinib exposure is not cytotoxic for GBM6 EGFRvIII specimens. We also identify a role of extracellular HA (via CD44) in altering the effect of erlotinib in GBM EGFR+ cells by modifying STAT3 phosphorylation status. Taken together, we report an in vitro tissue engineered platform to monitor signaling associated with poor response to targeted inhibitors in GBM.

## Introduction

Glioblastoma is the most common, aggressive, and deadly form of brain cancer.[1] Unlike many solid tumors where mortality is linked to metastases,[2] GBM mortality is driven by rapid diffusive spreading of GBM cells from the tumor margins throughout the brain along structural elements such as blood vessels.[3-6] The current standard-of-care is surgical resection (debulking) of the primary tumor mass followed by radiation and chemotherapy. Unlike surgical removal of many tumors where wide margins or total resection of the surrounding tissue is possible, debulking involves setting sharply defined surgical margins that cannot capture the diffuse physiological margin. GBM tumors recur rapidly (median: 6.9 mo post debulking) and at a site almost always in close proximity (>90% within 2 cm) of the original resection despite aggressive radiation and chemotherapy.[7] These GBM cells at the margins and those spreading into the parenchyma are the population that must be targeted therapeutically. Among GBM tumors, around 30% present a constitutively activated mutation of the EGF receptor (EGFRvIII) that presents a possible target for anti-tumor drugs.[8] EGFRvIII results from a truncation of the extracellular ligand-binding domain, and normally appears in conjunction with EGFR overexpression.

While targeting EGFR in GBM patients seems promising, in practice, current strategies are poorly effective in part due to both intrinsic and acquired resistance pathways, though mechanisms are still poorly understood.[9] Erlotinib is a reversible tyrosine kinase inhibitor (TKI) of EGFR, that binds to the intracellular TK domain of the receptor [10, 11] and was initially approved in 2004 as therapy for non-small cell lung cancer (NSCLC). EGFR activation needs both the association with an appropriate ligand and receptor dimerization on the cell surface, leading to phosphorylation of the intracellular domain of EGFR then activation of the PI3K/AKT and RAS/ERK signaling pathways. EGFR is amplified or overexpressed in many tumors, and EGFR TKIs have demonstrated better anti-tumor properties than chemotherapy in patients with aberrant EGFR, but typically TKI efficacy diminishes over a period of months.[12] Primary mechanisms that may contribute to erlotinib resistance include activation of alternative signaling, i.e. IGF1R pathway activation can influence EGFR inhibitor resistance via AKT regulation.[13] EGFRvIII GBM cancer cells may also alter PDGFRβ activation to maintain proliferation in response to EGFR inhibitors.[14] Another tactic in EGFR inhibitor resistance is activation of the PI3K/AKT/mTOR pathway. Mutations of PI3K-AKT pathway are present in 90% of GBM tumors,[15] are often accompanied by PTEN loss, and lead to the over-activation of the mTORC1 pathway.[16] Mutations in, or loss of, PTEN expression may be used as a marker of primary resistance to erlotinib. Additionally, STAT3, a transcription factor that regulates a variety of genes related to cell function and immune response to cancer may also influence tumor cells survival and resistance to apoptotic signals.[17] In some cases, EGFR inhibitors are able to inhibit phosphorylation of STAT3, thus preventing activation.[18]

In addition to acquired resistance, other factors may contribute to the failure of targeted therapies. The heterogeneity in the cellular constituents of EGFR-amplified GBM tumors, particularly the presence of GSCs and microglia, may contribute to higher intrinsic resistance to chemotherapeutic drugs and the capacity to recover from damage.[19, 20] Increasingly, it is believed that signals received from the tumor extracellular matrix may also contribute to acquired resistance,[21, 22] leading our effort to demonstrate tissue engineering platforms to investigate processes associated with EGFR inhibitor resistance. In support of these studies we have previously developed an adaptable gelatin hydrogel system to explore the role of brain-mimetic hyaluronic acid content in patient-derived GBM cell response to erlotinib.[23] HA has a dynamic role in the tumor microenvironment, both in its soluble and matrix-bound form.[24, 25] The objective of this work is to employ this tissue engineering platform to examine the combined effects of extracellular matrix signals from the GBM tumor microenvironment on response to repeated exposure to erlotinib. This effort requires three-dimensional hydrogel systems that recapitulate features of brain extracellular matrix as well as patient-derived GBM specimens [26] to provide cellular heterogeneity as well as established benchmarks of performance in xenograft models. Herein we explore the role of prolonged treatment cycles and extracellular HA on signal transduction pathway activation and resultant signature of resistance in EGFR+ vs. EGFRvIII GBM specimens.

## Materials and Methods

### Cell laden hydrogel fabrication

Gelatin methacrylamide (GelMA) and hyaluronic acid methacrylate (HAMA) were prepared as described previously.[27] We produced GelMA scaffolds that contained 0 or 1wt% HAMA in PBS (Invitrogen) by UV photopolymerization in the presence of 0.05 wt% LAP (Lithium phenyl-2,4,6-trimethylbenzoylphosphinate) as photoinitiator. Prepolymer solution (7 wt%) was pipetted into Teflon molds (5 mm diameter × 1.5 mm thick) and exposed to 10 mW/cm^2^ UV light (LED 365 nm) for 60 s. Patient-derived GBM6 and GBM12 cells were cultured within these hydrogels at a concentration of 4 million cells/ml. GBM6 and GBM12 specimens demonstrate different EGFR mutation status, clinical response to erlotinib, and degree of invasion in a patient-derived xenograft model (**Table 1**). EGFRvIII (GBM6) does not contain a ligand-binding domain and is constitutively active. GBM6 expresses both EGFR and EGFRvIII. Compression modulus (*E*) was determined via an Instron 5493 (100 N load cell) mechanical testing apparatus (20% strain/min). Diffusion coefficient (*D*) was calculated using fluorescence recovery after photobleaching (FRAP) assay, in combination with a FITC-dextran (40 KDa, Sigma-Aldrich) fluorescent probe.[27]

**Table 1.** Characteristics of GBM12 and GBM6 tumors. Fraction of glioma stem cells (GSC) and macrophage (Mϕ) / microglia were obtained by flow cytometry using CD133^+^ and CD68 cell markers. EGFR/PTEN; Primary/recurrent; Invasion rate (0: low, 7: high); Erlotinib efficacy in orthotopic mouse model, percentage of change in mean survival between control and erlotinib treatment groups (ref 36); MGMT (methylated /unmethylated); Subtype (neural, proneural, classical, mesenchymal); GSC (CD133^+^); Mϕ, microglia (CD68)

| GBM | EGFR/<br>PTEN | 1°/rec. | Invasion | erl % | MGMT | Subtype | GSC<br>% | Mϕ, microglia<br>% |
| --- | --- | --- | --- | --- | --- | --- | --- | --- |
| 12 | + / wt | 1° | 2 | 21 | M | M | 8.3 | 25.2 |
| 6 | vIII / wt | 1° | 6 | 5 | U | C | 38.0 | 17.0 |

### Cell culture and analysis of cell proliferation

GBM patient-derived xenograft (PDX) cells were cultured in DMEM medium supplemented with 10% FBS and penicillin/streptomycin (100 U/ml and 100 μg/ml). We used FBS media to explicitly examine the role of matrix microenvironment as a selection pressure, versus the study of the population dynamics of stem cell systems. Cells were cultured at 37°C in a 5% CO_2_ environment and used upon receipt from Mayo Clinic (passage 1). PDX cell laden hydrogels were incubated on an orbital shaker in low adhesion well plates containing standard culture media for 48 hours. After providing 48 hours to stabilize within the hydrogels, cells were exposed to a *1*^*st*^ *Dose* of 10 μM dose of erlotinib (Biovision Inc., in 0.1% v/v DMSO) added to the culture media for 3 days (0.1% v/v DMSO as a control). The erlotinib solution was subsequently replaced with fresh media for four day of recovery (days 4 – 7 of the culture), followed by exposure to a 10 μM second dose of erlotinib for an additional three days (2^nd^ dose; days 7 – 10 of the culture). The erlotinib concentration was selected as appropriate from previous experiments.[23] GBM-seeded hydrogel specimens were evaluated at the start and end of each erlotinib dose (**Scheme 1**). Total cell number per construct was determined using the PicoGreen DNA assay according to the manufacturer’s instructions (P11496; Invitrogen-Molecular Probes); sample ?uorescence at 520 nm (ex. 480 nm, Biotek Synergy HTX plate reader) was compared with DNA standards included in the assay kit. For some experiments, cells were exposed to the specific STAT3 inhibitor Stattic (SelleckChem; 5 μM) in combination with erlotinib (10 μM) for 3 days; metabolic activity was analyzed using the Vybrant® MTT cell proliferation assay kit (ThermoFisher Scientific). STAT3 inhibition was confirmed by Western Blot (**Figure S2**).

**Scheme 1.**
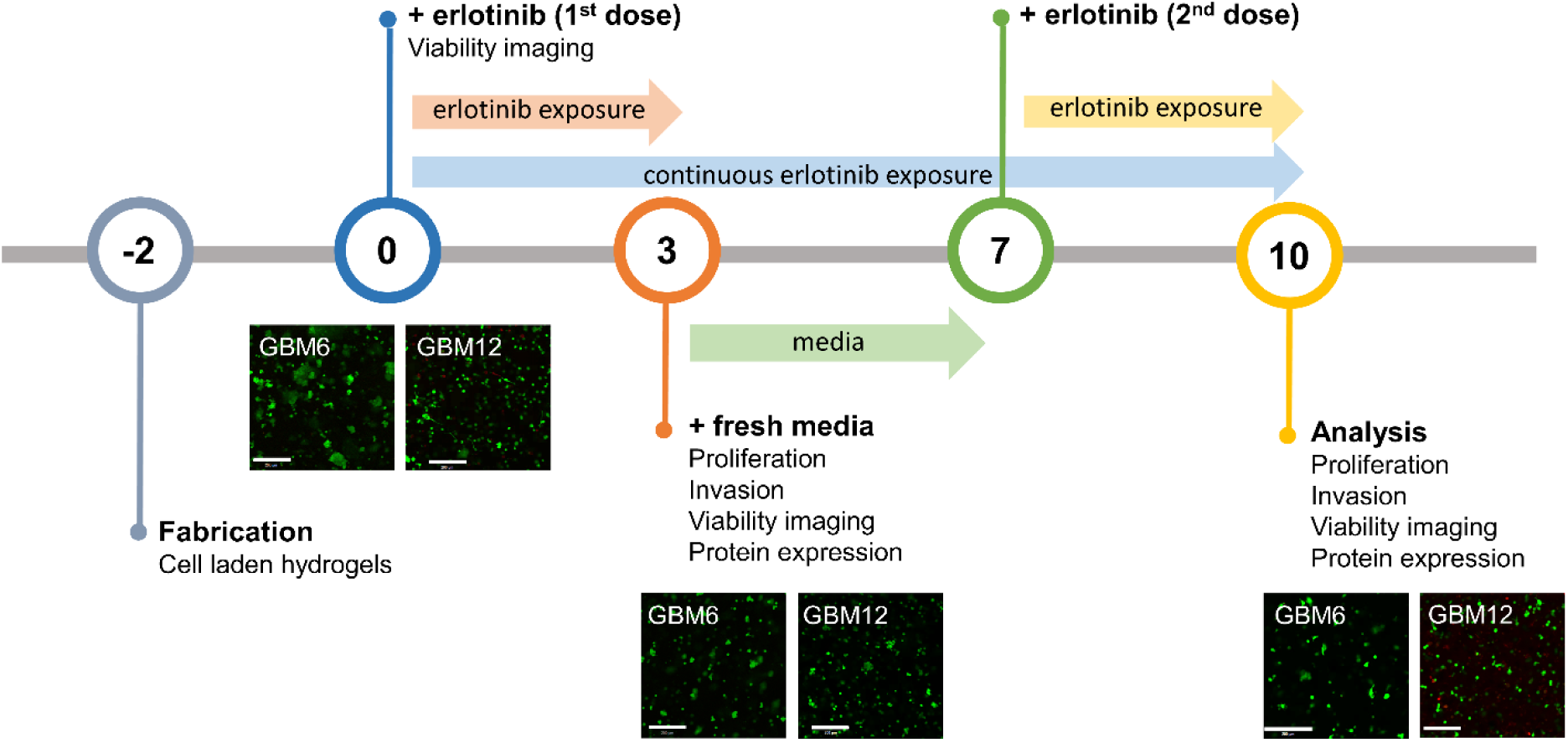
Treatment scheme of erlotinib administration (10 μM) to GBM6 and GBM12 within gelatin / HA hydrogels. Cells were exposed to erlotinib for 3 days (*1*^*st*^ dose), 2 days after encapsulation. Then erlotinib was eliminated from cell media for 4 days (fresh media), after that cells were exposed to a *2*^*nd*^ dose of erlotinib for 3 days. Cells were analyzed at days 0, 3, 7 and 10. Images show live/dead analyses on GBM cells within GelMA hydrogels.

### Ethics statement

The xenografts used in this study were established with tumor tissue from patients undergoing surgical treatment at the Mayo Clinic, Rochester, Minnesota. The Mayo Clinic Institutional Review Board approved these studies and only samples from patients who had provided prior consent for use of their tissues in research were included. All xenograft therapy evaluations were done using an orthotopic tumor model for glioblastoma on a protocol approved by the Mayo Institutional Animal Care and Use Committee.

### Analyzing cellular population using flow cytometry

The analysis of the cellular composition of PDX samples GBM6 and GBM12 focused on two subsets of cells: a CD133+ glioblastoma stem cell like cell population and a CD68+ tumor-associated microglia/macrophage (TAM) population[28]. Patient-derived GBM6 and GBM12 cells were stained with human-CD133 (PE conjugated 1:11 dilution; Miltenyi Biotec), then with human-CD68 (FITC conjugated 1:20 dilution; Invitrogen), following manufacturer’s protocol. A single-stained control of each dye and an unstained sample were also analyzed. Samples were stained with propidium iodide (PI, 1:1000, Thermo Fisher Scientific) for dead-cell exclusions. Flow cytometry was then performed using a BD LSR II flow cytometer (BD Biosciences). Acquired data were analyzed using FCS Express 5.0, and positive fraction of each cell type was obtained by subtracting the unstained control fraction from sample fraction.

### Quantifying cell invasion using a spheroid-based assay

Cell invasion was quantified for GBM spheroids embedded into the hydrogel matrix using a previously described method.[29] Briefly, 96 well-plates were coated with methylcellulose at 37°C overnight, forming a non-adherent gel surface. GBM cells were seeded into each well (500 cells/µL; ThermoFisher) overnight with constant horizontal-shaking (60 rpm) to form spheroids (37°C, 5% CO_2_). Individual spheroids (10K cells in size) were hand-picked via 200/1000 μl pipette tips, then placed into the Teflon mold with the hydrogel pre-polymer solution (30 μL/well, 7 wt% GelMA or 6 wt% GelMA+1wt% HAMA) followed by UV polymerization (10 mW/cm^2^; 60s). Hydrogel samples were then placed in regular media for 48 hours (day 0) prior to adding the *1*^*st*^ *Dose* of erlotinib (10 μM erlotinib vs. 0.1% control DMSO) for 72 hours. As with prior samples, the erlotinib was removed with samples placed in fresh media for four days followed by a *2*^*nd*^ *Dose* (10 μM erlotinib vs. 0.1% control DMSO) for an additional 3 days. Images of spheroids (n=6) were acquired on days 0, 3, 7 and 10 using a Leica DMI 400B florescence microscope under bright field, with cell invasion into the surrounding hydrogel matrix quantified via ImageJ as distance from the spheroid surface.[29]

### Protein immunoblotting

Analysis of protein expression was determined from cell-hydrogel specimens first lysed using RIPA buffer on ice for 30 min.[23] After lysis and determination of protein concentration, samples were separated on a 10% polyacrylamide precast electrophoresis gel (Biorad # 456-1096), and transferred onto a nitrocellulose membrane (Amersham Protan, GE Healthcare, # 10600012). Proteins were visualized by ECL western blotting detection reagents (Pierce, ThermoFisher Scientific), according to manufacturer’s instruction. The following primary antibodies were used (all rabbit; all from Cell Signaling, Danvers, MA): anti-β-actin (#4967); PDGF Receptor β (#4564); TORC1/CRTC1 (#2587); Phospho-p44/42 MAPK (Erk1/2) (#9101); p44/42 MAPK (Erk1/2) (#9102); PDGF Receptor α (#3164); Phospho-Stat3 (Tyr705) (#9131); Stat3 (#12640). Primary antibodies were then labeled with goat-anti-rabbit IgG conjugated to horseradish peroxidase (Cell Signaling). Blocking solution was 5wt% non-fat dry milk in TBST (Tris Buffered Saline with Tween® 20).

### Statistical analysis

All analyses were performed using a one-way analysis of variance (ANOVA) followed by Tukey’s HSD post-hoc test. Significance level was set at p < 0.05. At least n = 3 samples were examined for cell proliferation assays, at least n = 6 samples for invasion assays, and at least n = 3 samples were examined for Western Blot analysis. Error was reported in figures as the standard deviation unless otherwise noted.

## Results and discussion

### Cellular heterogeneity in tumor microenvironment

EGFR TKIs such as erlotinib reversibly bind to the intracellular catalytic TK domain of EGFR or EGFRvIII to inhibit auto-phosphorylation of the receptor as well as downstream signaling. While erlotinib shows a diverse variety of antitumoral responses versus GBM (reduced invasiveness and viability),[10] it is ineffective clinically. Here we explore the influence of cell and matrix properties of the tumor microenvironment on poor treatment response. GBM tumors contains gradients in matrix and oxygen content, varies temporally with disease progression, and differs significantly patient-to-patient.[30-34] GBM tumors contain a heterogeneous mix of clonal cells, microglia and a subpopulation of tumor initiating cells (glioblastoma stem cells, GSCs).[31, 35, 36] Under chemo- or radiotherapy, some clones become resistant and survive, increasing diversity and contributing to tumor malignancy and resistance to therapy even further.[37]

An opportunity to uncover new combinatorial therapies in GBM lies in defining potential axes of resistance and downstream targets. The potential of cancer cells to dynamically adjust their signaling to avoid TKIs complicates this mission. We have studied tumor cell heterogeneity in GBM6 and GBM12 glioblastoma lines (**Table 1**), focusing in stem cell and macrophage / microglia fractions. Signaling from EGFRvIII cells is constitutively active while EGFR+ could be constitutive or ligand activated.[10] Despite unsuccessful efforts trying to inhibit EGFR in GBM, such as the vaccine against EGFRvIII,[38] some tumors with particular characteristics have been responsive to TKIs.[39] Both GBM6 and GBM12 show molecular features (EGFR status and PTENwt) that make them theoretically susceptible to treatment with EGFR inhibitors (**Table 1**), yet GBM6 shows a significantly poorer response to erlotinib in vivo.[40] In this context, the development of ex vivo tumor platforms can provide insight regarding acquired resistance as a function of matrix environment as well as drug dosage and scheduling in a reduced time scale. We have previously developed an in vitro 3D tumor model that recreates key features of the tumor environment, notably the presentation of brain-mimetic hyaluronic acid,[27, 41] and we have shown that when used in combination with PDX, these hydrogels can offer a platform for profiling drug response.[23, 42] The inclusion of hyaluronic acid does not significantly alter either hydrogel mechanical properties (*E*_GelMA_ = 15.3 ± 2.1 and *E*_GelMA/HAMA_ = 17.5 ± 1.1 kPa) or diffusion coefficient (*D*_GelMA_ = 105.5 ± 7.1 and *D*_GelMA/HAMA_ = 104.2 ± 7.3 μm^2^/s). Therefore, we consider the presence of HA does not cause a significant architectural alteration of the microenvironment. However, since variations do exist, we are reporting data within experimental groups as relative to the DMSO control.

### Erlotinib inhibits proliferation and migration of EGFR+ GBM cells subject to matrix composition

We previously showed the significance of incorporating matrix-immobilized HA within gelatin hydrogels on GBM invasion capacity,[43] gene expression patterns[42] and the formation of robust endothelial cell networks as models of the tumor perivascular niche.[44] Therefore, we hypothesized that the presence of extracellular HA will affect the efficacy of cyclic erlotinib exposure. GBM cell-laden hydrogels were exposed to 10 μM erlotinib for 3 days, allowed to recover in erlotinib-free media for 4 days, then re-exposed to 10 μM erlotinib for an additional 3 days (to day 10). Cell response was monitored at day 3 (*1*^*st*^ *Dose*) and day 10 (*2*^*nd*^ *Dose*). Overall, GBM EGFRvIII-expressing cells maintain growth and invasion potential in spite of repeated erlotinib doses (**Figure 1**). However, we observed significantly reduced EGFR+ GBM12 proliferation (p<0.05) after the *1*^*st*^ *Dose* of erlotinib exposure in both gelatin and HA-modified gelatin hydrogel. While we observed significant reduction in invasion after erlotinib exposure in gelatin-only hydrogels, GBM invasion was not reduced with erlotinib in the presence of matrix-immobilized hyaluronic acid. Notably, GBM6 vIII cells were only sensitive to erlotinib in hydrogel environments containing HA (21% reduced proliferation p<0.05 compared to control DMSO), while invasion remained unchanged (representative image of GBM6 invasion, **Figure 1c**). Multiple rounds of erlotinib treatment led to a greater decrease of cell number (60%, p<0.05 control) and invasion (67% p<0.05 control) of GBM12 and proliferation of GBM6 (38%, p<0.05 control) tumor cells in gelatin-only hydrogels. Interestingly, inclusion of brain-mimetic hyaluronic acid blunted the effect of repeated erlotinib exposure, with extended erlotinib treatment not significantly reducing GBM cell proliferation or migration behavior. This finding reinforces the need to create defined extracellular matrix environments to study the dynamic response of GBM cells to EGFR inhibitors.[45, 46] Together, these findings suggest inhibition of proliferation and migration by erlotinib is influenced by matrix composition, subject to the presence of vIII mutation, with proliferation findings consistent with previously reported mouse model results.[40]

**Figure 1.**
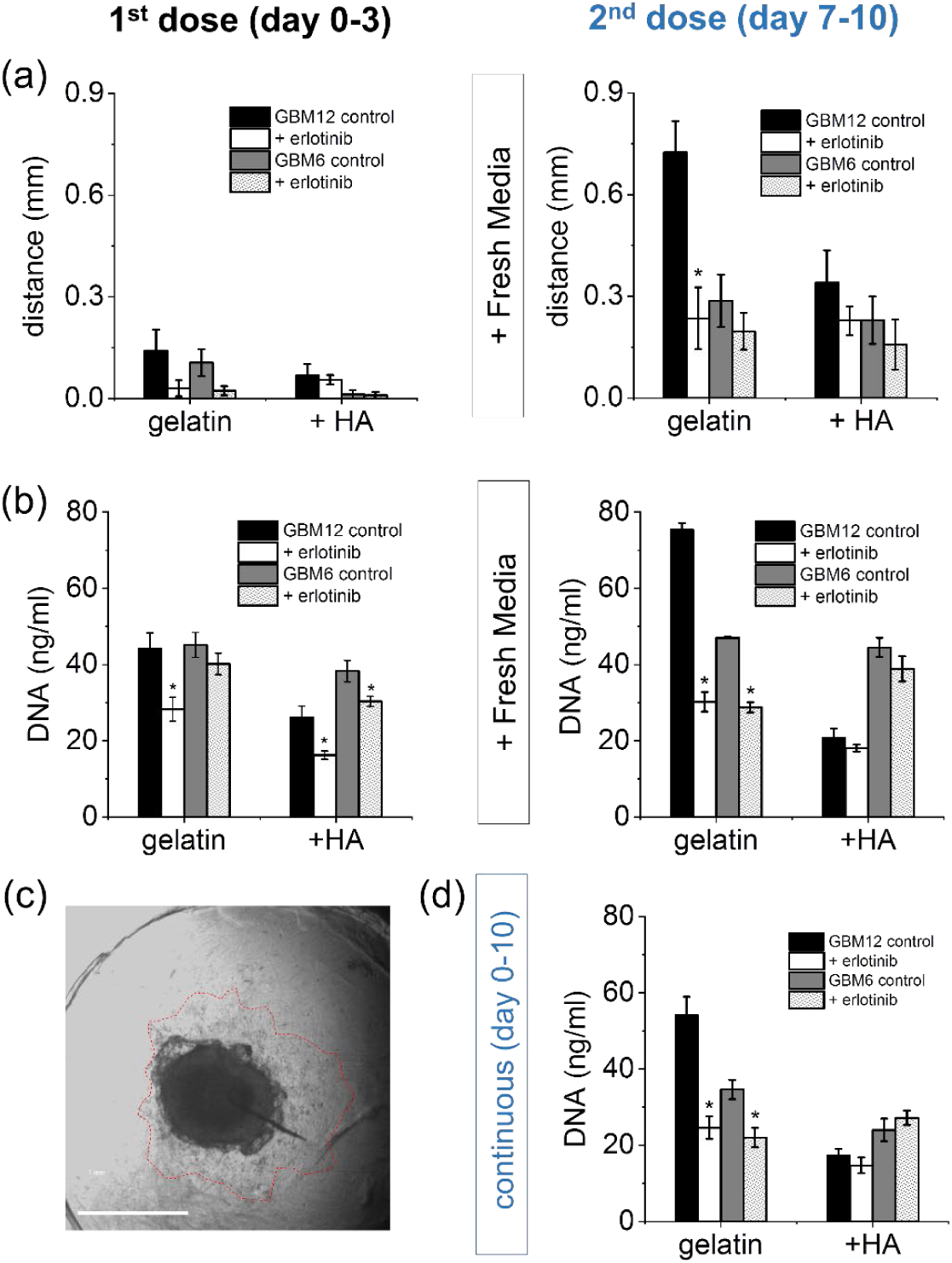
(a) Quantification of GBM cell invasion into the GelMA based hydrogel when exposed to erlotinib, as compared to DMSO control (0.1% v/v). Cells are exposed to 10 µM erlotinib dose twice from day 0 to day 3 and from day 7 to day 10. From day 3 to day 7, cell are culture in fresh DMEM media (10% FBS). Erlotinib in GelMA hydrogels containing 1wt% HA showed more limited inhibition of invasion, especially in GBM6 vIII. GBM12 EGFR+ cells in GelMA hydrogels showed significantly reduced invasion after second dose of erlotinib. *p < 0.05 compared to DMSO control. (b) DNA quantification (PicoGreen) of GBM cells in GelMA and GelMA / 1wt% HAMA hydrogels. Both GBM6 and GBM12 cell proliferation is inhibited by erlotinib, however, cell number is not affected in HA matrices after the second dose (day 10). (c) Invasion of GBM6 cells in GelMA hydrogels at day 7. (d) Similar results are obtained when cells are exposed to erlotinib continuously throughout the 10 day period. *p < 0.05 compared to DMSO control.

Alternatively, cells were exposed to erlotinib continuously for the 10 day period (media was changed at days 3 and 7, comparable to cyclic experiments) to assess potential differences between dosing schedule (cyclic vs. continuous). The dosing schedule is an important parameter in treatment success[47, 48] and can contribute resistance.[49] In general, continuous dosing is still the most common clinical strategy for delaying tumor growth and invasion as compared to pulsatile administration of high dose TKIs.[50] We observed no differences between discontinuous and continuous treatment of tumor platforms with 10 μM erlotinib in terms of overall number of viable cells within the hydrogels (**Figure 1d**).

### Activation of proliferation and survival pathways in response to erlotinib

We subsequently examined expression patterns for key components of EGFR signal transduction pathway in GBM[8] (ERK, mTOR, STAT3) as a result of exposure to multiple discontinuous doses of erlotinib. Initially, we looked at ERK expression, since this pathway is important in cancer cell survival, and resistance of vIII cells to erlotinib has been suggested to be related to upregulated PI3K.[40] Recent studies suggest that intrinsic resistance to erlotinib in GBM EGFR+ tumors is related to a rapid adaptive response provoked by an increase in TNF signaling[51] that reactivates ERK. After exposure to erlotinib, ERK phosphorylation decreases in both xenograft samples, more significantly in GBM6 vIII, regardless of cyclic dosing status or matrix composition (**Figure 2**). However, this pathway does not seem to be dominant in the observed differences in proliferation and migration (**Figure 1**). The influence of erlotinib on the Ras/ERK pathway is also more notable in vIII cells, with EGFR+ GBM12 only becoming sensitive after a second dose.

**Figure 2.**
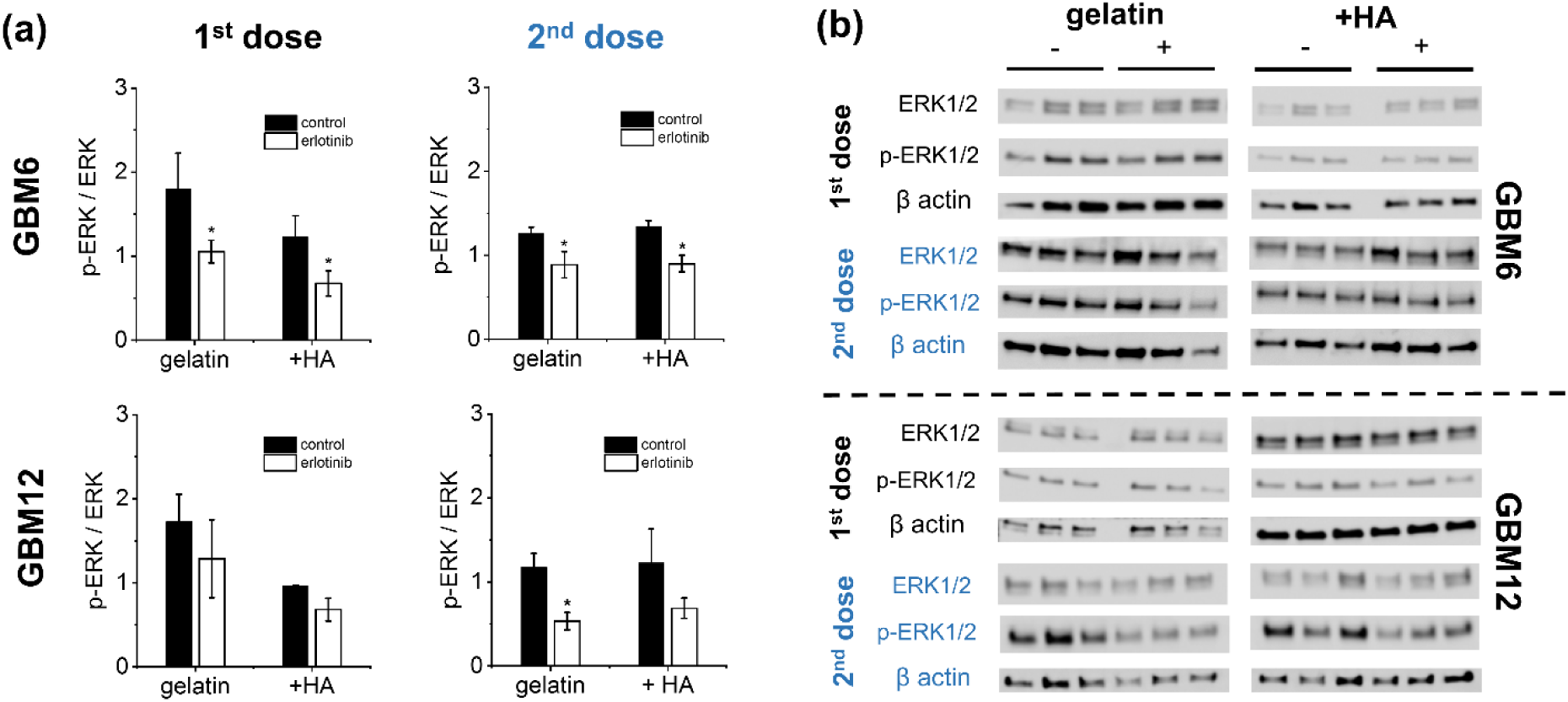
EGFR inhibition with 10 µM erlotinib induces downregulation of ERK in GBM6 and GBM12 PDX cells. DMSO is used as control (0.1% v/v). (a) Quantification of p-ERK1/2 calculated from Western blot with the indicated antibodies (b) in GelMA and GelMA / HA hydrogels. Results normalized against a β-actin loading control. Concentration of p-ERK decreases in all gelatin-only GBM cultures exposed to second dose of erlotinib (day 10), while only GBM6 seem to be responsive to first dose (day 3). *p < 0.05. Data are presented as mean ± s.e.m, n=3.

We subsequently explored the role of the PI3K / Akt / mTOR pathway via mTORC1 protein expression (**Figure 3**), finding this pathway less sensitive to initial and repeated erlotinib treatments, than ERK1/2, in both cell types. PTEN acts as a negative regulator of PI3K, although this protein is commonly inactivated in glioblastoma, both GBM6 and GBM12 present the wild type PTEN. Surprisingly, the PI3K / Akt / mTOR pathway was upregulated only in GBM6 vIII cells after a second dose of erlotinib, observations frequently related to acquired resistance phenotypes (**Figure 3**).

**Figure 3.**
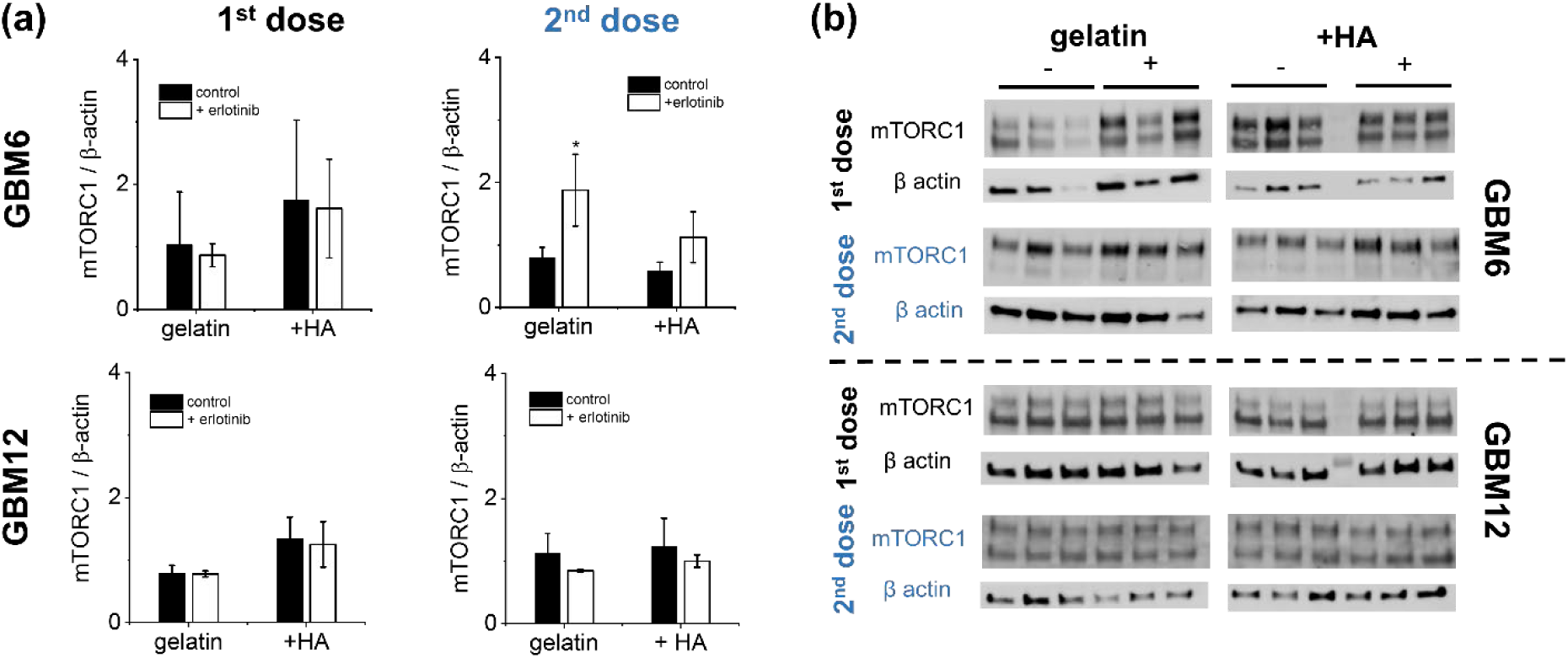
EGFR inhibition with 10 µM erlotinib does not affect PI3K / mTORC1 signaling in GBM6 and GBM12 PDX cells. DMSO is used as control (0.1% v/v). (a) Quantification of mTORC1 calculated from Western blot with the indicated antibodies (b) in GelMA and GelMA / HA hydrogels. Results normalized against a β-actin loading control. Concentration of mTORC1 is not affected by short (day 3) or long (day 10) term EGFR inhibition. *p < 0.05. Data are presented as mean ± s.e.m.

These observations do not coincide with shifts in PI3K phosphorylation (**Figure S1**) that remains unchanged in GBM6. This could be explained by alternative activation of mTORC1 through PKC and independently of PI3K / Akt, associated with low efficacy of therapies targeting these proteins in glioblastoma.[52]

Signal transducer and activator of transcription 3 (STAT3) is a transcription factor that regulates expression of genes related to cell cycle and survival in GBM.[53] In normal cells, levels of activated STAT3 are transitory. However, STAT3 remains constitutively active in the majority of solid tumors, including glioblastoma.[54] Phosphorylation of the transcription factor STAT3 is significantly increased on the activation site in EGFRwt and EGFRvIII-expressing tumors.[45] To evaluate the activation of JAK / STAT3 pathway in response to erlotinib exposure, we examined STAT3 protein expression (**Figure 4**). GBM12 shows some downregulation in STAT3 activation, but only in hydrogel environments lacking brain-mimetic HA. Long-term cyclic exposure to erlotinib increases STAT3 phosphorylation in GBM6 vIII cells, but not in GBM12 EGFR+, consistent with reports that associate EGFRvIII to regulation of STAT3.[53]

**Figure 4.**
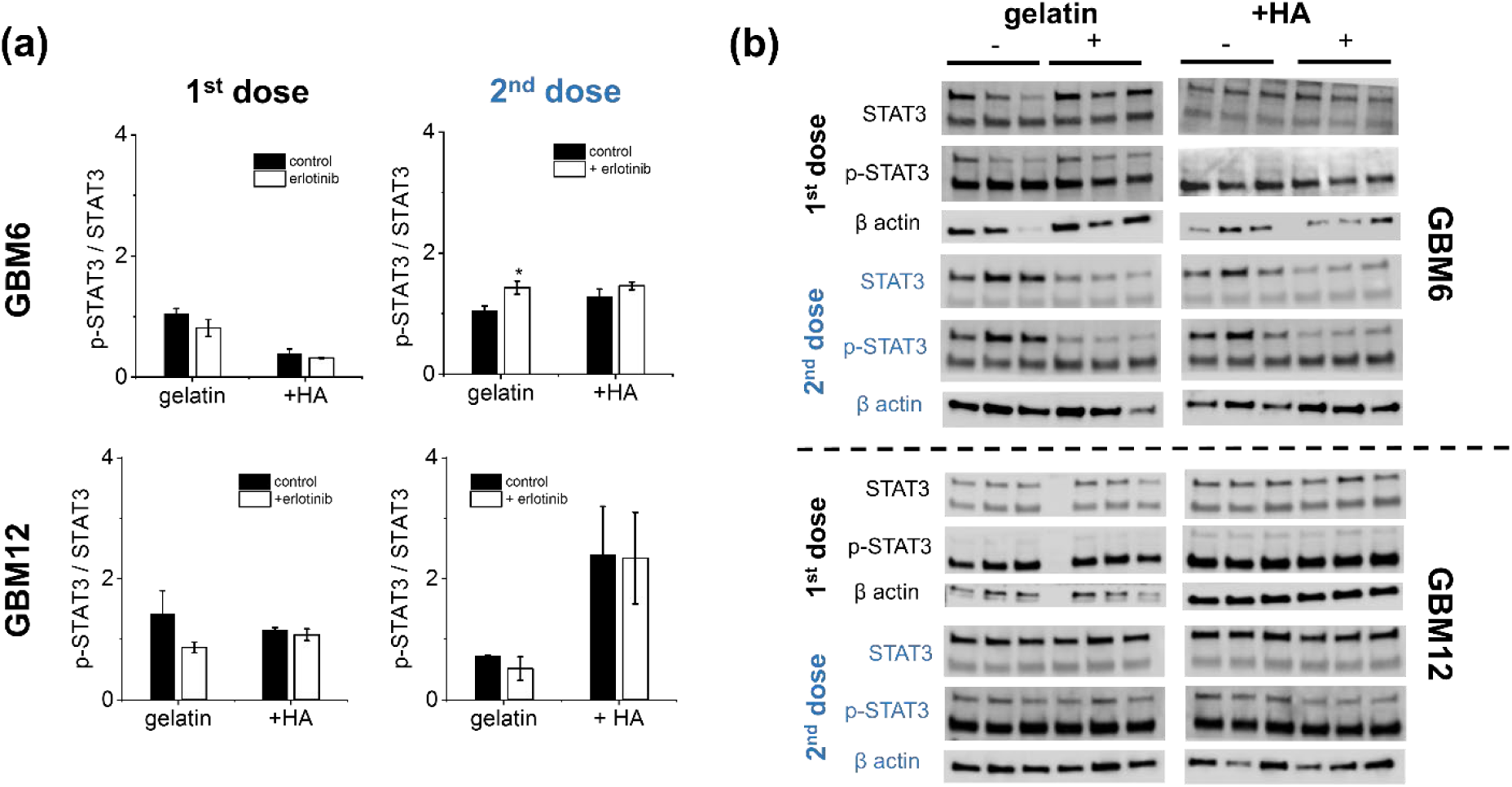
Extracellular HA affects EGFR inhibition with 10 µM erlotinib through STAT3 signaling in GBM6 and GBM12 PDX cells. DMSO is used as control (0.1% v/v). (a) Quantification of p-STAT3 calculated from Western blot with the indicated antibodies (b) in GelMA and GelMA / HA hydrogels. Results normalized against a β-actin loading control. *p < 0.05. Data are presented as mean ± s.e.m.

Hyaluronic acid represents a significant portion of the brain extracellular matrix. In addition to providing a local physical structure, HA provides a wide range of biophysical signals to GBM tumor cells. In other solid tumors, activation of STAT3 has been associated to hyaluronan deposition,[55] but no data exists about the relationship of the HA matrix environment in GBM influencing signaling response to drug exposure. We observe p-STAT3 is upregulated in EGFR+ GBM12 cells, but only in matrices containing brain-mimetic HA and only after repeated erlotinib exposure. This increase in STAT3 activity could be associated with the lack of efficacy of erlotinib in HA matrices, as observed in proliferation and invasion profiles (**Figure 1**).

### Activation of compensatory pathways: Erlotinib promotes upregulation of PDGFRβ in GBM6 vIII cells

GBM tumors often exhibit PDGF autocrine signaling not present in normal brain tissues, and have been shown to express both surface receptors (PDGFRα and PDGFRβ).[56] PDGFRα is the second most frequently amplified RTK in GBM behind EGFR, and their protein co-expression occurs in 37% of GBM.[42] Indeed, the lack of EGFR / PDGFRα co-expression in GBM tumors is related to reduced efficacy of TKIs.[23] We observed GBM6 vIII tumors express reduced levels of PDGFRα, regardless of matrix composition and number of doses (**Figure 5**). However, PDGFR levels are not altered in EGFR+ GBM12 regardless of matrix composition or exposure to erlotinib. It has been reported that EGFR signaling negatively regulates PDGFRβ transcription, via mTORC1, in glioblastoma models.[14] Interestingly, our study reveals that repeated inhibition of EGFR in GBM6 vIII cells via erlotinib upregulates PDGFRβ and is associated with upregulation of mTORC1, especially in gelatin-only matrices (**Figure 3**). Taken together, these data suggest that HA-rich matrices do not seem to provide signals to preferentially stimulate PDGFR expression patterns versus gelatin platforms in the course of the erlotinib treatment.

**Figure 5.**
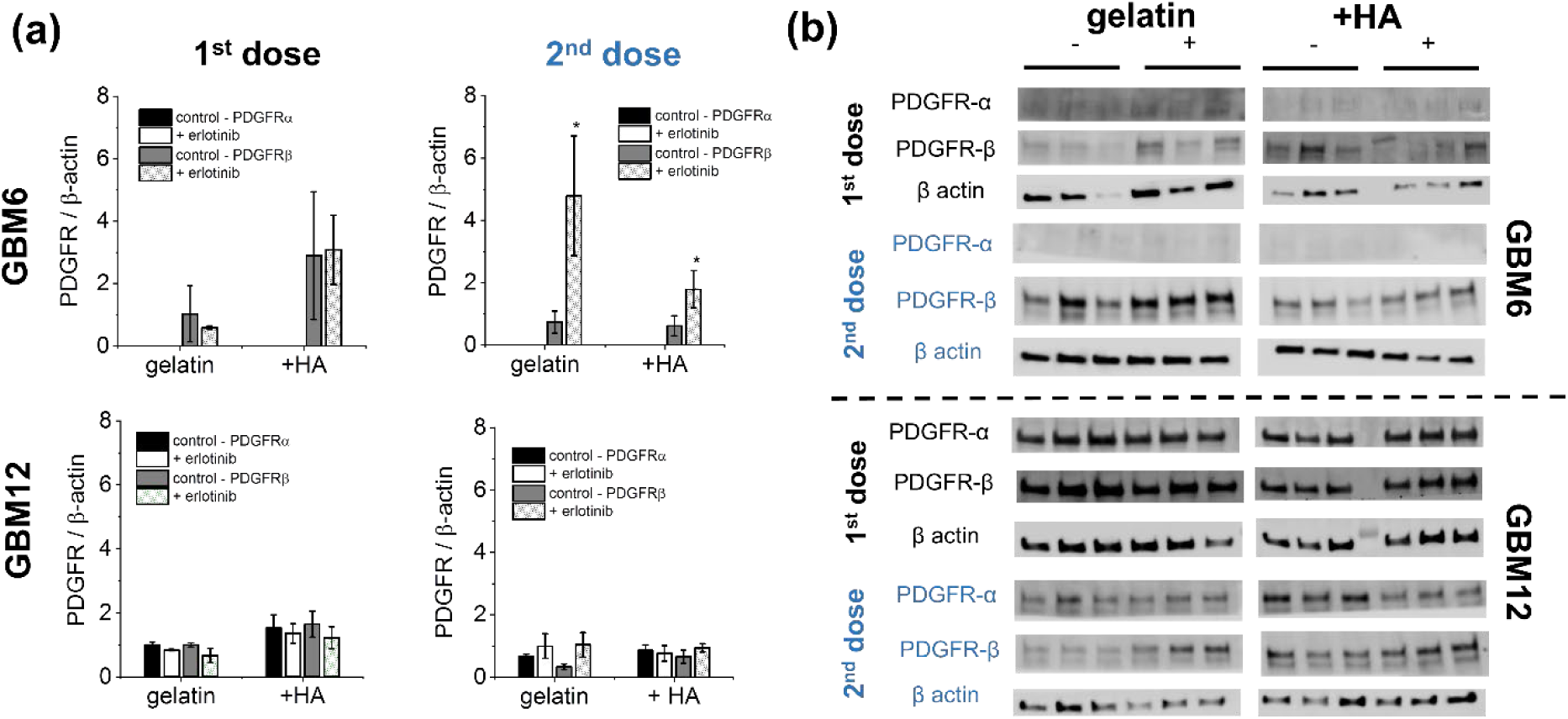
EGFR inhibition with 10 µM erlotinib upregulates the expression of RTK PDGFRβ in GBM6 vIII cells after second dose. DMSO is used as control (0.1% v/v). (a) Quantification of PDGFR was calculated from Western blot with the indicated antibodies (b) in GelMA and GelMA / HA hydrogels. Results normalized against a β-actin loading control. *p < 0.05. Data are presented as mean ± s.e.m.

### Influence of extracellular matrix microenvironment in erlotinib efficacy: Role of hyaluronic acid through CD44 signaling

We next examined whether matrix-bound HA within the hydrogel environment affects how xenograft variants respond to multiple rounds of erlotinib exposure. While EGFR+ cells (GBM12) appear particularly sensitized to erlotinib in gelatin-only hydrogels (**Figure 1**), the effect is lost in HA-decorated gelatin hydrogels after repeated erlotinib doses. We also observed no significant differences in proliferation for EGFRvIII GBM6 cells exposed to erlotinib in HA-decorated gelatin hydrogels. Erlotinib was also not effective inhibiting invasion in HA-decorated gelatin hydrogels for both GBM specimens. These signatures of resistance could be explained by activation of alternative signaling pathways (e.g. CD44-EGFR activation) that lead to mTORC1 – STAT3 activation and subsequent HA synthesis. This loop could then promote increased cell activity when erlotinib is added a second time, including to an extent within GelMA-only hydrogels.

To investigate more deeply the potential hallmarks of TKI resistance as a function of matrix biophysical signals, we compared the phosphorylation levels of STAT and ERK when the HA receptor CD44 was blocked (Anti-CD44 Rat mAb (A020), EMD Millipore, 3 μg/ml) in conjunction with erlotinib exposure (**Figure 6**). In EGFR+ GBM12, activation of ERK decreases upon CD44 blockade in gelatin matrices, while p-STAT is downregulated in HA-containing matrices only when both EGFR and CD44 are inhibited, via erlotinib and α-CD44, respectively. However, CD44 engagement does not seem to alter EGFR signaling response in GBM6 vIII, as p-STAT3 and p-ERK are not deactivated upon addition of αCD44. Moreover, the inhibition of STAT3 activation with Stattic sensitizes GBM12 cells to erlotinib in gelatin matrices containing HA (**Figure 7**). These results corroborate the importance of STAT3 in a successful inhibitory effect of erlotinib in glioblastoma, in particular in extracellular activation of EGFR-CD44 by HA in GBM12 EGFR+ tumors (**Figure 8**).

**Figure 6.**
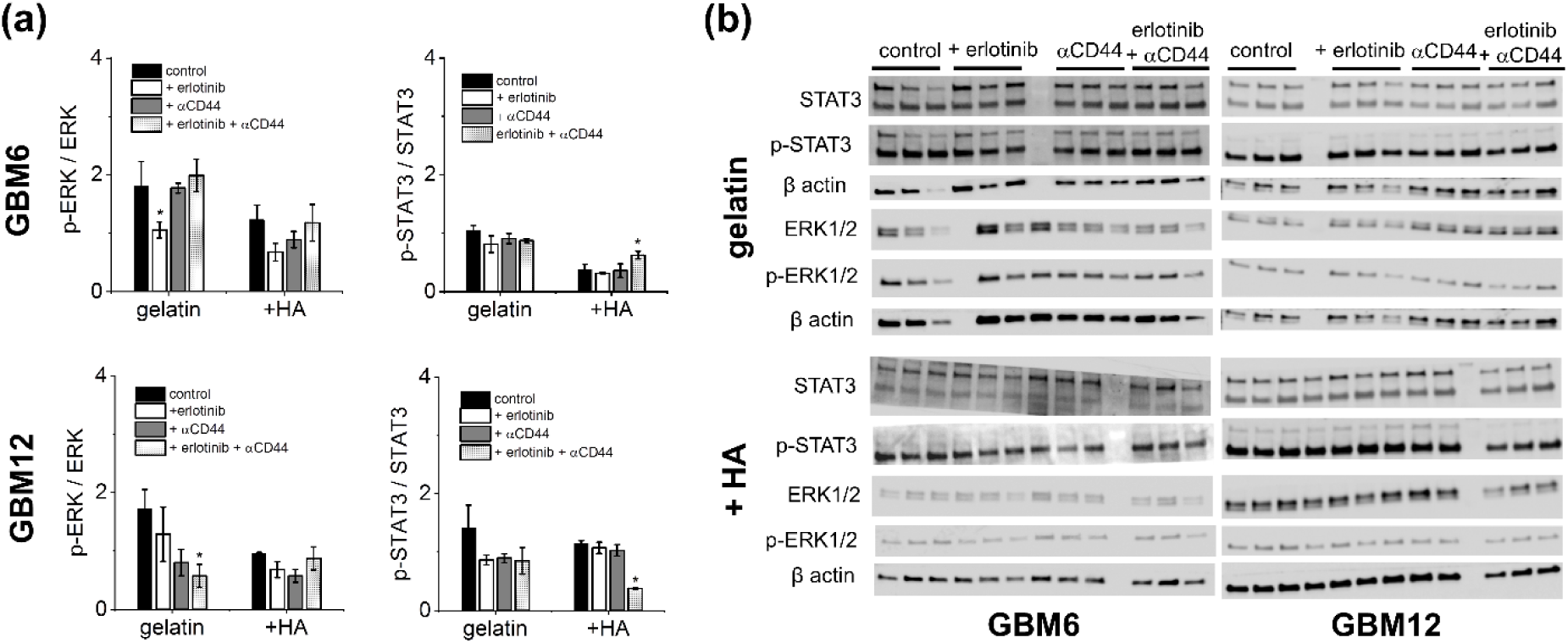
Combinatorial inhibition of EGFR and CD44 with 10 µM erlotinib and ?CD44 (3 µg/ml) downregulates p-STAT3 in GBM12 EGFR+ cell platforms as opposed to GBM6 vIII PDX cells, showing the regulatory effect of CD44 in EGFR+ cells when HA is present in ECM. DMSO is used as control (0.1% v/v). (a) Quantification of p-STAT and p-ERK was calculated from Western blot with the indicated antibodies (b) in GelMA and GelMA / HA hydrogels. Results normalized against a β-actin loading control. *p < 0.05. Data are presented as mean ± s.e.m.

**Figure 7.**
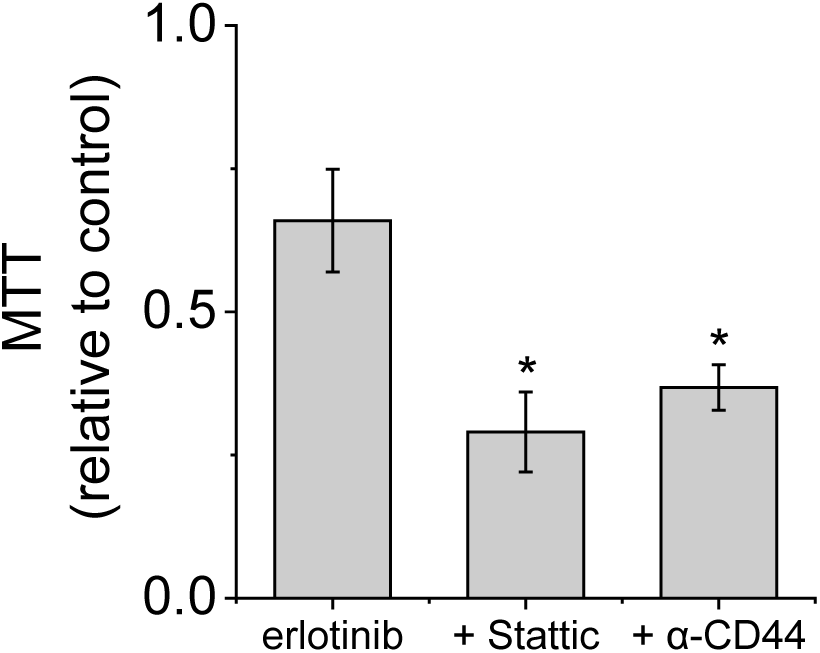
Inhibition of STAT3, in combination with erlotinib, reduces metabolic activity (MTT) of GBM12 PDX cells in GelMA/HAMA hydrogels. Cells are exposed to 10 μM erlotinib and 5 μM Stattic dose for 3 days. DMSO is used as control (0.1% v/v). *p < 0.01. Data are presented as mean ± s.e.m, n=3.

**Figure 8.**
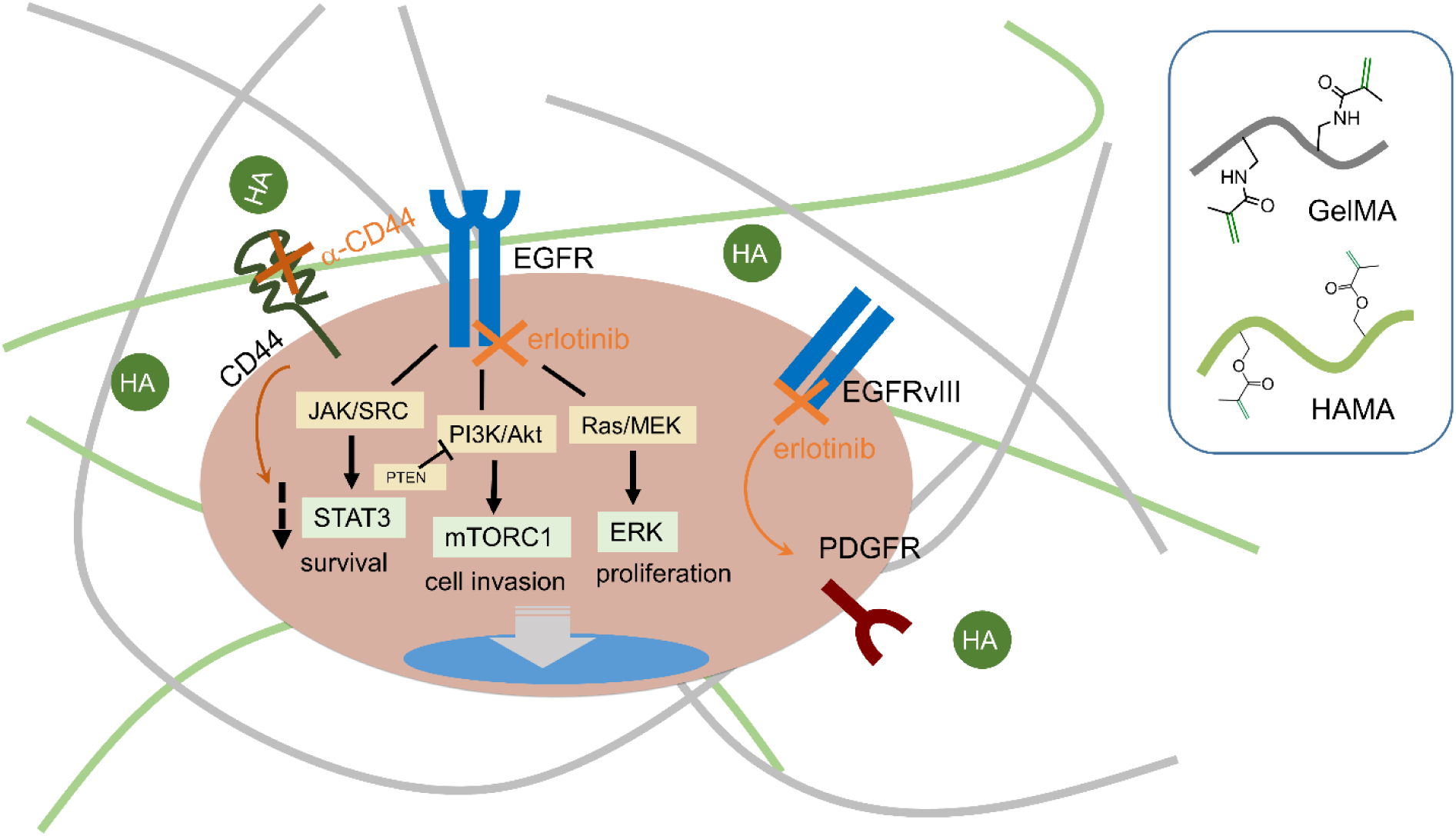
Influence of extracellular HA in erlotinib efficacy using a reductionist 3D glioblastoma model. The epidermal growth factor receptor (EGFR) signaling pathway regulates growth, survival and proliferation in GBM through three main signaling pathways: JAK/STAT, PI3K/mTOR and Ras/ERK. The addition of erlotinib, together with the inhibition of extracellular HA receptor (CD44), downregulates STAT3 phosphorylation, only in EGFR+ (GBM12) cells. HA microenvironment also decreases erlotinib faculty to stop cell motility in these cells, compared to control, but does not influence ERK phosphorylation. PDGFRβ expression is enhanced in GBM6 vIII, upon EGFR inhibition.

## Conclusions

Resistance to anticancer drugs is a complex process that arises from a variety of dynamic factors such as the role of the immune system, behavior of neighboring cells, tumor tissue physiology, mutations and alterations in signaling pathways. Improved understanding of acquired resistance may provide new insights into the biology of EGFR-influenced GBM tumors. We report analysis of mechanisms commonly associated with erlotinib resistance utilizing a 3D hydrogel platform that recreates features of the glioblastoma microenvironment. Notably, this ex vivo platform allows for a rapid dissection of mechanistic insights commonly associated with therapeutic resistance. We show that repeated dosing of erlotinib promotes expression of PDGFRβ in GBM6 vIII cells and highlight the importance of extracellular hyaluronic acid in a favorable inhibition of EGFR, through STAT3 deactivation. Gelatin and hyaluronic acid act as a structural network within the brain, but we show here matrix-bound hyaluronic acid has an active role in the efficacy of targeted therapies. As the presence of hyaluronic acid in the tumor microenvironment is constantly changing via cell mediated hyaluronic acid-remodeling and synthesis, these findings motivate future studies to more precisely elucidate the role of extracellular hyaluronic acid (matrix bound vs. soluble; role of molecular weight and remodeling) in therapeutic efficacy.

## Acknowledgements

The authors would like to acknowledge the members of the Roy J. Carver Biotechnology Center at the University of Illinois Urbana-Champaign for their advice and assistance with experiments for this manuscript. Specifically, the authors would like to thank Barbara Pilas in the Flow Cytometry Facility. Research reported in this publication was also supported by the National Cancer Institute (R01CA197488, BACH), National Institute of Diabetes and Digestive and Kidney Diseases (R01 DK099528, BACH), and the National Institute of Biomedical Imaging and Bioengineering (T32EB019944, JEC) of the National Institutes of Health. The content is solely the responsibility of the authors and does not necessarily represent the official views of the NIH. The authors are also grateful for additional funding provided by the Department of Chemical & Biomolecular Engineering and the Carl R. Woese Institute for Genomic Biology at the University of Illinois at Urbana-Champaign.

## Data availability

All data analyzed during this study are included in this published article (and its supplementary information file). Other raw data required to reproduce these findings are available from the corresponding author on reasonable request.

