## Supplementary information for "Hyaluronic acid-functionalized gelatin hydrogels reveal extracellular matrix signals temper the efficacy of erlotinib against patient-derived glioblastoma specimens"

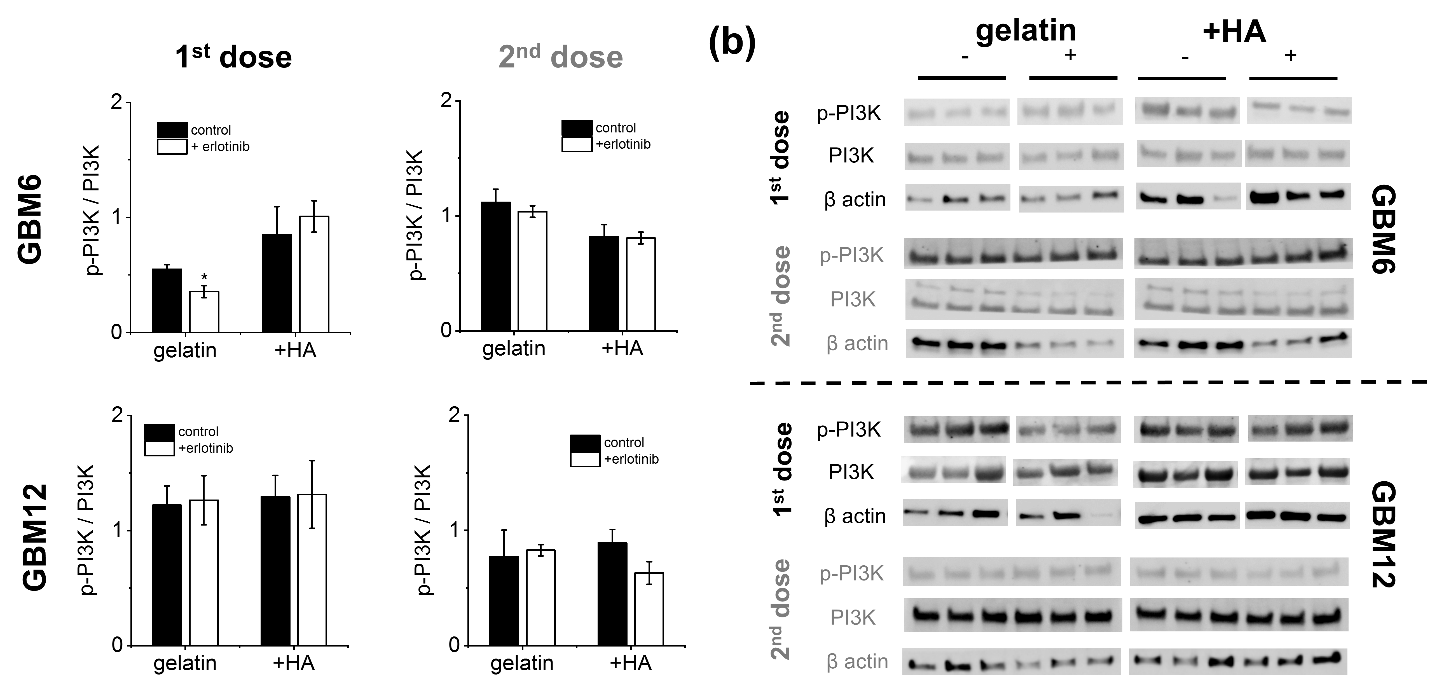

**Figure S1.** EGFR inhibition with 10 μM erlotinib does not modify phosphorylation of PI3K in GBM6 vIII and GBM12 EGFR+ PDX cells regardless of dosing or matrix composition. DMSO is used as control (0.1% v/v). (a) Quantification of p-PI3K was calculated from Western blot with the indicated antibodies (b) in GelMA and GelMA / HA hydrogels. Results normalized against a β-actin loading control. *p < 0.05. Data are presented as mean ± s.e.m. Data of p-PI3K after 1^st^ dose are extracted from a previous publication (ref. 23). We are adding them here to highlight the trend of the newly presented data for the second dose.

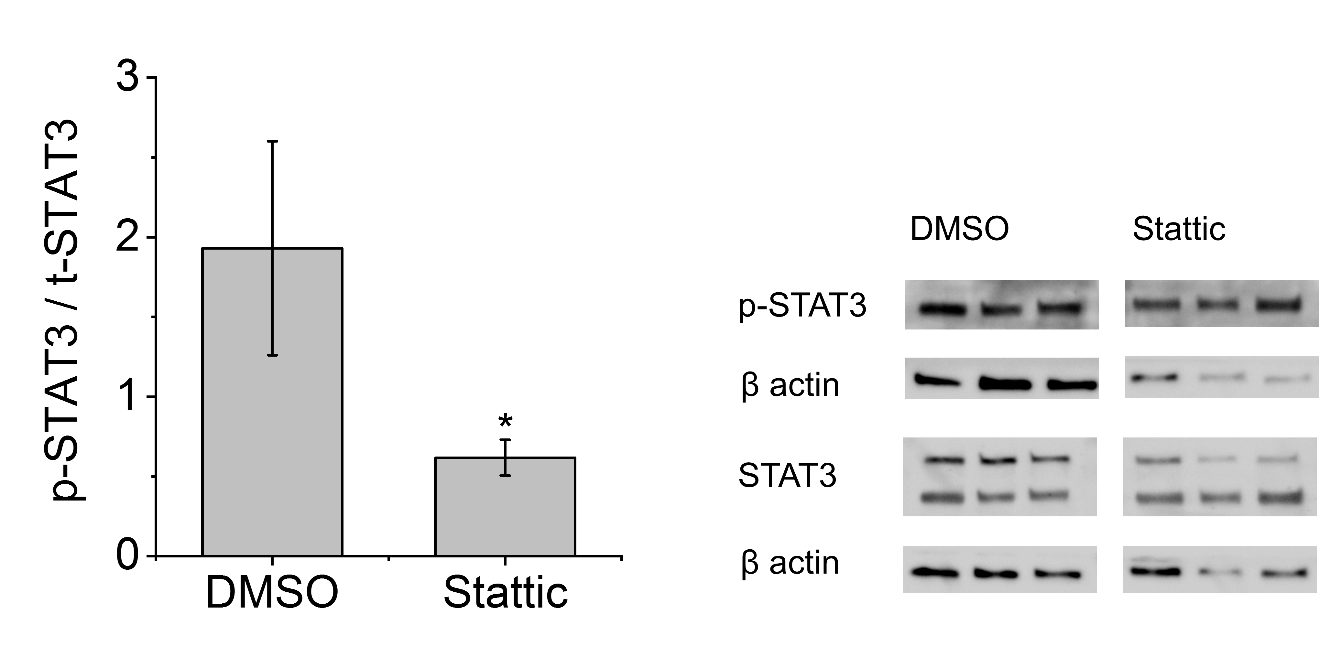

**Figure S2.** Inhibition of STAT3 phosphorylation in GBM12 EGFR^+^ cells in GelMA / HAMA hydrogels via addition of STAT3 inhibitor Stattic (5 μM). Control: 0.1%v/v DMSO. The phosphorylated:total STAT3 level is reported from Western blot, with results normalized against a β-actin loading control. *p < 0.05. Data are presented as mean ± s.e.m, n = 3.
